# Glycoprotein-glycoprotein receptor binding detection using bioluminescence resonance energy transfer (BRET)

**DOI:** 10.1101/2023.01.21.525003

**Authors:** Kamila Adamczuk, Adolfo Rivero-Müller

## Abstract

The glycoprotein receptors, members of the large G protein-coupled receptors (GPCRs) family, are characterized by a large extracellular domains responsible of binding their glycoprotein hormones. Hormone-receptor interactions are traditionally analyzed by ligand-binding assays most often using radiolabeling but also by thermal shift assays. However, the use of radioisotopes requires appropriate laboratory conditions, and moreover, for this purpose, purified cell membranes are most often used instead of living cells. This in turn poses another challenge due to the altered stability of membrane proteins in detergents used for purification. Here, we overcome such limitations by applying bioluminescence resonance energy transfer (BRET) in living cells to determine hormone-receptor interactions between a *Gaussia* luciferase (Gluc) luteinizing hormone/chorionic gonadotropin receptor (LHCGR) fusion and its ligands (yoked human chorionic gonadotropin (yhCG) or luteinizing hormone (LH)) fused to the enhanced green fluorescent protein (eGFP). We first show that the Gluc-LHCGR is expressed on the plasma membrane and is fully functional, as well as the chimeric eGFP-hormones that are properly secreted and able to bind and activate the WT LHCGR. Finally, we applied the method to determine the interactions between clinically relevant mutations in the hormone as well as the receptor and show that this assay is fast and effective, plus safer and cost efficient alternative to radioligand-based assays, to screen for mutations in either the receptor or ligand. It enables kinetic measurements in living cells, detection of biosynthesis of the receptor (membrane expression) and it is compatible with downstream cellular assays - including firefly luciferase-based readouts.

## Introduction

G protein-coupled receptors (GPCRs) are the most numerous group of membrane receptors responsible for the transduction of extracellular signals into the cell in response to external stimuli in the form of neurotransmitters, hormones, growth factors or light ^1^. They are distinguished by the presence of a seven α-helix transmembrane domain connected by three extracellular and three intracellular loops. Due to the number and wide variety of receptors belonging to this family, they play key roles in physiological processes, including the nervous, endocrine, reproductive and cardiovascular systems ^2^. Therefore, they are the target of around 30% of commercial drugs ^3^. GPCRs include receptors for glycoprotein hormones among which is the luteinizing hormone/chorionic gonadotropin receptor (LHCGR), involved in the development of secondary sexual characteristics and synthesis of progesterone in females and testosterone in males ^4^. Due to the occurrence of polymorphisms and mutations in the genes encoding both the LHCGR and its ligands, it is important to understand the impact of these alterations on the receptor-ligand functions.

Radioligand-binding assays have traditionally been the basic tools for studying the interactions between ligands (or agonist/antagonists) and GPCRs, which is of particular importance for the development of the pharmaceutical industry. These assays are based on the incubation of radioligand with membranes from cells, or very rarely whole cells, expressing the GPCR of interest followed by the measurement of radioactivity bound ligand ^5^. Hitherto, the most frequently used radiolabeled ligands for this purpose are ^3^H- and ^125^I-labeled ligands ^6^. The utility of assays based on radiolabeling is mainly due to their high sensitivity and the fact that radioisotopes only slightly modify the chemical structure of the ligand. Nonetheless, the major disadvantage of radioligand-binding assay is that the preparation of labeled ligands is hazardous^7^. Therefore, the radioligand-based assays require specific laboratory conditions and they are associated with the production of radioactive waste. Furthermore, some of the radioisotopes are characterized by their short half-lives ^6^. Radioligand binding assay is one of the endpoint assays, that measures receptor bound vs. unbound radio-ligand, usually on purified membranes, what precludes analyzing the kinetics of their interactions in living cells. Moreover, the use of membranes, often including the endoplasmic reticulum and Golgi apparatus, prevaricates obtaining information on the subcellular localization and activation of the receptor. Ligand-receptor interactions are also commonly analyzed using the thermal shift assay, which measures the thermal stability of the purified receptor in the presence and absence of ligand ^8^.

Due to the above-mentioned issues, assays using non-radioactive labels and living cells are sought after. In this case, it is important to ensure that the label (or tag) does not affect the ligand affinity to the receptor and does not disturb the interaction between the receptor and the ligand^9^. The most commonly used nonradioactive bioassays are based on the fluorescence resonance energy transfer (FRET) and bioluminescence resonance energy transfer (BRET) phenomena. In the case of FRET, two chromophores are used, one of which is excited resulting in energy transfer and excitation of the second chromophore and thus its fluorescence emission ^10^. FRET is detected by the change of fluorescent ratio between the donor and the acceptor, which equals to large background to noise ratio and variability, thus the need of measuring single cells ^11,12^ instead of populations, this is labor-intensive and time-consuming. In the case of the BRET, one fluorescent protein is used as an acceptor, while a bioluminescent enzyme acts as a donor. As a result of enzymatic oxidation of a substrate, the energy is released and transferred to the fluorescent acceptor. A necessary condition for both of these phenomena to occur is a small distance of <10nm (<100Å) between the donor and the acceptor enabling energy transfer ^13^. The BRET method differs from the FRET method in that it does not require excitation with a light source, nevertheless it requires the use of a substrate suitable for a chosen luciferase, thus it has lower background noise and variation.

To achieve BRET we engineered a novel donor:acceptor pair, this is the use of *Gaussia* luciferase (Gluc) and the enhanced green fluorescent protein (eGFP). The function of bioluminescent energy donor is performed by Gluc which is linked to the *N*-terminus of the LHCGR. Gluc is an enzyme naturally secreted by the copepod *Gaussia princeps*. It is responsible for the oxidative decarboxylation of coelenterazine without additional cofactors, resulting in the formation of coelenteramide and the emission of blue light (480nm) ^14,15^. Gluc is also one of the smallest known luciferases with a mass of 19.9 kDa and it is distinguished by the fact that the humanized form of Gluc generates an over 100-fold higher bioluminescent signal as compared to Firefly and Renilla luciferases ^16^. In turn, the function of the fluorescent energy acceptor here is performed by eGFP fused to both of the LHCGR ligands: human chorionic gonadotropin (hCG) or luteinizing hormone beta subunit (LHβ). The eGFP is characterized by the presence of two substitutions consisting of Ser65Thr and Phe64Leu, resulting in 35-fold brighter fluorescence as compared to the wild-type (WT) GFP ^17,18^. Most importantly, this fluorescent protein is excited at 488nm and thus it coincides with the emission peak of Gluc ^19^.

Here, we report a BRET system to study the interaction between the LHCGR and both of its ligands – hCG and LH. Furthermore, we show that this method can be used to study clinically relevant mutations both in the ligand and the receptor. In order to achieve that, we generated the only previously-described mutation on common glycoprotein alpha subunit (CGA) - Glu80Ala ^20^, as well as three mutations in the extracellular domain (ECD) of LHCGR: Cys131Arg ^21^ and Ile152Thr ^22^, reported as binding-deficient, and Glu354Lys reported as binding-capable but signal-deficient ^23^. The CGA subunit was selected due to the fact that it is the only one known mutation in the *CGA* gene reported so far. Most likely, the lack of other reports on mutations within this gene is due to the essential functions performed by CGA subunit - it is necessary for the formation and proper functioning of LH, follicle-stimulating hormone (FSH) and thyroid-stimulating hormone (TSH) as well as hCG, the latter is indispensable for implantation and maintenance of pregnancy ^24^. The CGA^Glu80Ala^ mutation was found in a patient with malignant neoplasm. Hitherto, no functional assays have been performed to demonstrate the importance of this mutation, in conjunction with hCGB/LHB, for its ability to bind and activate LHCGR ^20,24^.

## Materials and Methods

### Cell culture and transfection

All cell lines were cultured in Dulbecco’s Modified Eagle medium (DMEM)/F12 (Gibco) containing 10% fetal bovine serum (FBS), 100 μg/mL streptomycin and 100 IU/mL penicillin (Gibco) in CO2 incubator at 37°C and 95% humidity. Human embryonic kidney (HEK293) cells were obtained from ATCC, whereas HEK293 cells stably expressing a luminescent cAMP GloSensor (GS-293) (Promega) were previously created in our lab ^25^. Another cell line that has been used in this study is the GS-293 line stably transfected with the plasmid encoding the LHCGR and referred to as GS-LHCGR cell line ^26^. Transfections were performed using the Turbofect Transfection Reagent (ThermoFisher Scientific #R0531) according to the manufacturer’s protocol at 70-80% cell confluence.

### Molecular cloning

The plasmids used as templates for further cloning were yhCG, NLS-AmCyan-P2A-LHB^WT^-mCherry and LHCGR^WT^-P2A-mCherry, whereas mCherry-TOPO and pUC18 plasmids were used as negative mock transfection controls. The eGFP sequence derived from eGFP-NPM WT Addgene (Plasmid #17578) was cloned into the yhCG (a gift from Prema Narayan ^27^) vector at the carboxyl terminus, from here onwards referred to as yhCG^WT^-eGFP. In the case of the second ligand, the procedure was analogous and the eGFP coding sequence was introduced at the carboxyl terminus of the LHβ subunit ^26^. Furthermore, the NLS-AmCyan part was removed from the construct which was then named LHB^WT^-eGFP.

In turn, for the donor the sequence encoding Gluc derived from pCMV-*Gaussia* Luc Vector (ThermoFisher Scientific #16147) was introduced at the *N*-terminus. Several variants of Gluc-LHCGR differing in length of flexible domain as well as the presence of Gluc signal peptide or the presence of mCherry coding sequence were created. Gluc-LHCGRv1 is distinguished by the presence of Gluc signal peptide between the LHCGR signal peptide and LHCGR coding sequence as well as the presence of a linker composed of 10 amino acids (GGSGGGGSGG) between the sequences for Gluc and LHCGR. Similarly, Gluc-LHCGRv2 also contains a flexible domain built of 10 amino acids (GGSGGGGSGG), whereas the coding sequence for Gluc signal peptide was removed. The flexible domains of Gluc-LHCGRv3 and Gluc-LHCGRv4 are composed of 5 amino acids (GGGGS), nonetheless in the case of the latter, the mCherry coding sequence was also removed.

All cloning aside from the mCherry and NLS-AmCyan removal were performed using the Gibson Assembly. For this purpose, a linear vector and DNA insert with overlapping ends were obtained, which were then mixed together with Gibson Assembly Master Mix - Assembly (NEB #E2611) and incubated at 50°C for 15 minutes. Subsequent steps involve transformation of the obtained products into *E.coli* cells, purification of plasmids and their verification by Sanger sequencing (Genomed). On the contrary, mCherry and NLS-AmCyan sequences were removed from constructs using REPLACR-mutagenesis ^28^. The plasmids reported in this work can be obtained via Addgene https://www.addgene.org/Adolfo_Rivero-Muller/.

### Flow cytometry

HEK293 cells were seeded in 12-well plates in DMEM/F12 medium. On the following day, cells were transiently transfected with plasmids encoding either the wild-type LHCGR^WT^-P2A-mCherry or Gluc-LHCGR^WT^ clones. Furthermore, mCherry-TOPO plasmid was used as an additional control for transfection efficiency, while pUC18 plasmid was used as mock transfected negative control. 48 hours after transfection, cells were incubated first with primary antibody referred to as rabbit anti-HA tag antibody (Cell Signaling #3724S) at 37°C for 1h and subsequently with donkey anti-rabbit IgG (H+L) Highly Cross-Adsorbed Secondary AntibodyAlexa-647 (ThermoFisher Scientific #A-31573) at 37°C for 1h. The primary antibody was diluted in ratio 1:1600, whereas the secondary antibody was diluted in ratio 1:1000 and TBS containing 2% bovine serum albumin (BSA) was used for dilution. The fluorescence intensity at the cell surface was read using FACSCalibur flow cytometer (BD). Mean fluorescence intensity (MFI) was plotted to compare relative expression levels at the cell surface of LHCGR^WT^-P2A-mCherry and Gluc-LHCGR^WT^ clones.

### Imaging of living cells

HEK293 cells were seeded in 24-well imaging plate (MoBiTec #5231-20) and on the following day cells were transiently transfected with plasmids encoding the yhCG, yhCG^WT^-eGFP, NLS-AmCyan-P2A-LHB^WT^-mCherry and LHB^WT^-eGFP. In the case of plasmids encoding the LHβ subunit, cells were transfected either alone or co-transfected with CGA to generate a functional LH composed of both subunits. The mCherry-TOPO plasmid was used as negative mock transfected control and transfection efficiency control as previously. 48 hours upon transfection, cells were imaged using A1 Nikon Eclipse Ti confocal microscope with 640 nm laser.

### cAMP analysis

LHCGR activation was detected by cAMP generation using the GS-293 cell line, which was seeded in 96-well plates. On the next day, cells were transiently transfected with either the LHCGR^WT^-P2A-mCherry or Gluc-LHCGR^WT^ plasmids. 48h post transfection, DMEM-F12 medium was replaced with assay medium, which consisted of DMEM-F12 and CO_2_-independent medium (Gibco #18045088) in ratio 1:1. The assay medium was supplemented with 2% GloSensor Reagent (Promega #E1291) and 0.1% BSA. The following step was the equilibration of cells in assay medium for 1 hour at room temperature. Afterwards, the measurement of baseline luminescence was made for 15 min which was followed by the stimulation of GS-293 cell line with recombinant hCG (rec-hCG) or recombinant LH (rec-LH). cAMP production was measured as a luminescent readout with the use of Tecan M200Pro microplate reader. The second part of the experiment involved the use of GS-LHCGR cell line distinguished by stable expression of the LHCGR receptor. GS-LHCGR cells were seeded in 96-well plate, while at the same time HEK293 cells were seeded in 6-well plates. On the following day, HEK293 cells were transiently transfected with yhCG^WT^, yhCG^WT^-eGFP, NLS-AmCyan-P2A-LHB^WT^-mCherry and LHB^WT^-eGFP. In the case of plasmids encoding the LHβ subunit, cells were transfected either alone or co-transfected with a plasmid encoding CGA. 36 hours upon transfection DMEM-F12 medium was replaced with fresh cell culture medium. 48 hours post transfection, the cell culture medium from 6-well plates was collected and either concentrated with Amicon Ultra microconcentrator 10 kDa cut-off (Merck Millipore #UFC501096) or not. Meanwhile, DMEM-F12 medium in 96-well plates was replaced with assay medium, the cells were transferred to the microplate reader and incubated in assay medium for 1 hour at room temperature. Afterwards, the measurement of baseline luminescence was made as described above and both concentrated and non-concentrated media were used to stimulate GS-LHCGR cells. Activation of LHCGR was measured as described above.

### Fluorescence and luminescence measurements

Next step of the study was the fluorescence measurement of the collected and concentrated medium containing secreted hormones (yhCG, yhCG^WT^ -eGFP, NLS-AmCyan-P2A-LHB^WT^-mCherry/CGA and LHB^WT^-eGFP/CGA). 36 hours upon transfection DMEM-F12 medium was replaced with fresh DMEM/F-12 medium without phenol red (Gibco #21041025). 48 hours post transfection, the cell culture medium from 6-well plate was collected and either concentrated with Amicon Ultra microconcentrator 10 kDa cut-off (Merck Millipore #UFC501096) or not. Afterwards, fluorescence top reading was performed (exc. 488 nm, em. 509 nm). Furthermore, the luminescence measurement of medium collected from cells transfected with either the LHCGR^WT^-P2A-mCherry or Gluc-LHCGR^WT^ plasmids was performed. Both luminescence and fluorescence measurements were read using Tecan M200Pro microplate reader.

### Equalization of eGFP-fused hormones

To normalize the concentration of eGFP-coupled hormones secreted into the cell culture medium, the GloSensor Assay was performed using the GS-LHCGR cell line. First, HEK293 cells were seeded in 6-well plates, whereas GS-LHCGR cells were seeded in 96-well plates. At 70% confluency, HEK293 cells were transfected with plasmids encoding either yhCG^WT^, yhCG^WT^-eGFP or TOPO-mCherry as mock transfected control. Furthermore, cells were cotransfected using either NLS-AmCyan-P2A-LHB^WT^-mCherry and CGA or LHB^WT^-eGFP and CGA plasmids. 36 hours upon transfection cell culture medium in 6-well plates was replaced with fresh medium. 48 hours post transfection, the cell culture medium from 6-well plate was collected and equalized by fluorescence, using Amicon Ultra microconcentrator 10 kDa cut-off. Meanwhile, 11 different dilutions were prepared for both recombinant hormones - hCG and LH. Then, DMEM-F12 medium of the 96-well plates was replaced with assay medium, the cells were transferred to the microplate reader and incubated in assay medium for 1 hour at room temperature.

Afterwards, the measurement of baseline luminescence was made and GS-LHCGR cells were stimulated with either concentrated or non-concentrated cell culture medium as well as using different dilutions of recombinant hormones. Activation of LHCGR was measured as described above.

### BRET assay

HEK293 cells were seeded in 96-well plates and 6-well plates and on the following day cells were transiently transfected. In the case of 96-well plates, cells were transfected with either the WT LHCGR-P2A-mCherry or Gluc-LHCGRv4, whereas the other plate was transfected with either the WT yhCG^WT^, yhCG^WT^-eGFP or cotransfected with either NLS-AmCyan-P2A-LHB^WT^-mCherry and CGA or LHB^WT^-eGFP and CGA plasmids. 36 hours upon transfection DMEM-F12 medium in 6-well plates was replaced with fresh DMEM/F-12 medium without phenol red (Gibco #21041025). 48 hours post transfection, the cell culture medium was collected and concentrated as previously. Meanwhile, HEK293 cells in 96-well plates were washed with DPBS, then concentrated hormones were added to the wells and cells were incubated with hormones for 20 minutes at room temperature. After incubation, cells were washed twice with DPBS, then DPBS and coelenterazine (Promega #S2001) were added to the wells at a final concentration of 20 μM. Thereafter, the measurement of luminescence and fluorescence top reading (exc. 230 nm, em. 509 nm) were performed using Tecan M200Pro microplate reader.

### Application of ligand-binding assay based on BRET to investigate the CGA subunit mutation and LHCGR binding-deficient mutations

In order to demonstrate the functionality of the method described in this study, a two-part experiment was performed. First part of the experiment was carried out to assess whether the mutant hCG is able to bind and activate the LHCGR. To prepare the vector coding for yhCG^Glu80Ala^-eGFP, REPLACR mutagenesis was performed as described by Trehan et al^28^. Initially, GS-LHCGR cells were seeded in 96-well plates, while HEK293 cells were seeded in 6-well plates. On the following day, HEK293 cells were transiently transfected with either the yhCG^WT^-eGFP or yhCG^Glu80Ala^-eGFP plasmids. 48 hours after transfection, the cAMP measurement was carried out as described above. The next step was the imaging of living cells. For this purpose, HEK293 cells were seeded in 24-well imaging plate and on the following day cells were transiently transfected with plasmids encoding either the yhCG^WT^-eGFP or yhCG^Glu80Ala^-eGFP plasmids. 48 hours after transfection, cells were imaged as previously. BRET was analyzed as following: HEK293 cells were seeded in 96-well plates and 6-well plates. On the following day, cells seeded in 96-well plates were transiently transfected with Gluc-LHCGR^WT^, whereas the cells seeded in 6-well plates were transfected with either yhCG^WT^-eGFP or yhCG^Glu80Ala^-eGFP. 36 hours later, medium from 6-well plates was replaced with fresh DMEM-F-12 medium without phenol red and 48 hours upon transfection, the cell culture medium was collected and concentrated as above. Then, the BRET experiment was performed as described in the BRET assay section.

The second part of the experiment was carried out to assess the effect of Cys131Arg, Ile152Thr and Glu354Lys LHCGR mutants on their ability to bind ligand and transduce signal. REPLACR mutagenesis was performed as described before ^28^. As a result, we obtained plasmids referred to as Gluc-LHCGR^Cys131Arg^, Gluc-LHCGR^Ile152Thr^ and Gluc-LHCGR^Glu354Lys^. Then, the membrane expression of the resulting constructs was assessed by flow cytometry. For this purpose, HEK293 cells were seeded in 12-well plates in DMEM/F12 medium and on the next day they were transiently transfected with plasmids encoding either Gluc-LHCGR^WT^ or mutant receptors. 48 hours after transfection, the flow cytometry analysis was performed as described above.

### Sequential measurements of ligand-receptor (BRET) interactions and receptor activity

GS-293 cells were seeded in 96-well plate, whereas HEK293 cells were seeded in 6-well plate. On the following day, GS-293 cells were transiently transfected with either the LHCGR^WT^-P2A-mCherry or Gluc-LHCGR^WT^ plasmids, while HEK293 cells were transfected with either the LHB^WT^-eGFP/CGA or NLS-AmCyan-P2A-LHB^WT^-mCherry/CGA plasmids. 36 hours later, medium from 6-well plate was replaced with fresh DMEM/F-12 medium without phenol red and 48 hours upon transfection, the cell culture medium was collected and concentrated as above. 48h post transfection, HEK293 cells were equilibrated in GloSensor assay medium for 1 hour at room temperature, then the measurement of baseline of Firefly luciferase (FFluc) luminescence (cAMP) was made for 15 minutes, followed by stimulation with LHB^WT^-eGFP/CGA. Then a readout was performed for another 45 minutes in a plate reader. After completion, cells were incubated for 1 hour before addition of hormones, wash twice with DPBS, and addition of coelenterazine, followed by the measurement of luminescence and fluorescence.

### Statistical analysis

GraphPad Prism 9 software (Graph Pad Software, San Diego, CA, USA) was used for statistical analysis using one-way ANOVA.

## Results

A representation of the of the constructs (**Figure 1A**) and the principle - where BRET is used to detect the interaction (binding) of the ligand, either LH or hCG, tagged with eGFP, and the LHCGR, fused with Gluc (**Figure 1B**). Several different architectures were used to ensure the proper biosynthesis, localization and functionality of both the ligands and the receptor.

**Figure 1.**
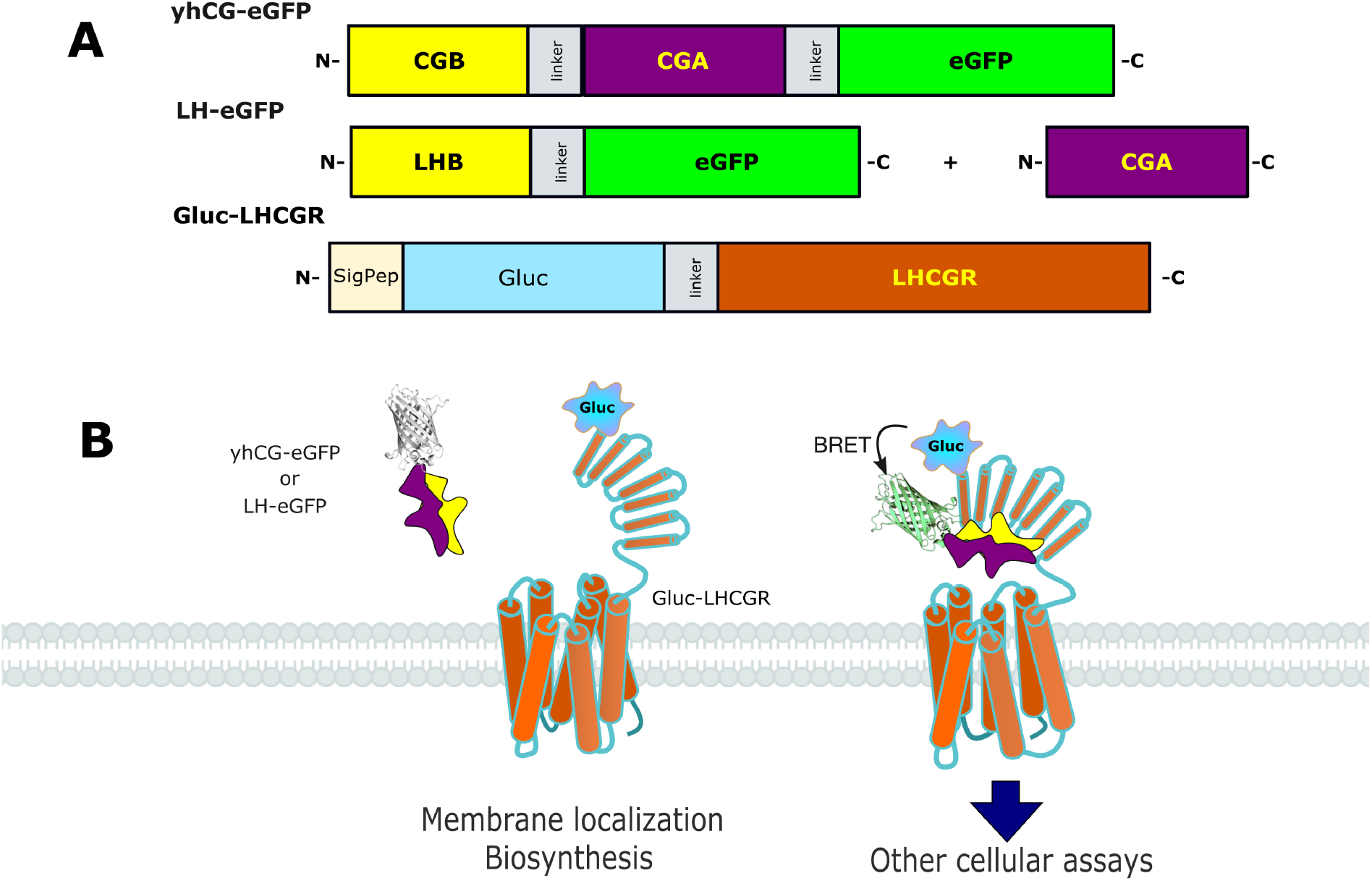
Schematic representation of the BRET-based glycoprotein-LHCGR assay. **(A)** Representation of the genetic constructs: the acceptor, a fusion of yhCG or LHB to eGFP via a short linker. The donor, in turn, was created by the fusion of Gluc to the extracellular domain of the LHCGR, where luciferase is preceded by the signal peptide of the LHCGR (SigPep). Gluc and LHCGR are also joined by a protein linker to allow flexibility. **(B)** Schematic representation of the resulting proteins and their cellular localization, as well as the resulting BRET phenomenon upon interactions between the acceptor and donor in the presence of coelenterazine – Gluc’s substrate.

### Membrane expression of the receptor

The first stage of the study was to assess the membrane expression of Gluc-LHCGR with different architectures using flow cytometry. While the majority of Gluc-LHCGR clones resulted in no membrane expression, Gluc-LHCGRv4 was distinguished by a high membrane expression (**Figure 2A**), which was 1.23-fold higher than that the wild-type (WT) LHCGR.

**Figure 2.**
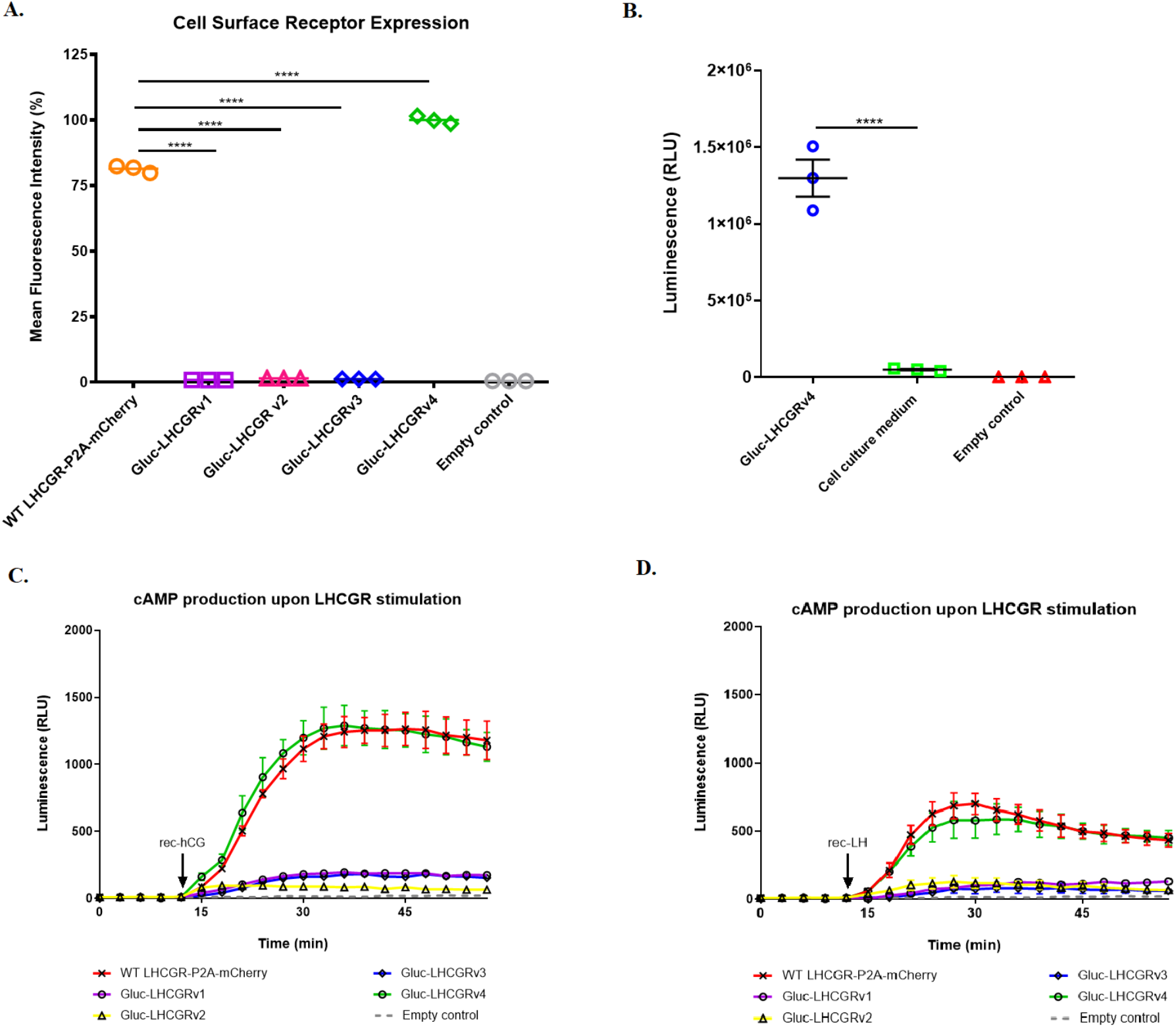
Molecular characterization of Gluc-LHCGR variants. **(A)** Flow cytometric analysis showed that the membrane expressions of Gluc-LHCGRv1, Gluc-LHCGRv2 and Gluc-LHCGRv3 were negligible and thus comparable to the expression of mock transfected control cells. On the contrary, the percentage median fluorescence intensity (MFI) of the Gluc-LHCGRv4 was 1.23-fold higher as that of the LHCGR^WT^-P2A-mCherry. Data is expressed as MFI ± standard error of the mean (SEM) of three independent experiments. **** p<0.0001. **(B)** Luminescence measurement carried out directly in wells with cells expressing Gluc-LHCGRv4 revealed that the luminescence was over 26-fold higher than the luminescence observed in the medium taken from abovementioned wells. Data is representative of an experiment performed in triplicate and was repeated independently at least three times. **** p<0.0001 **(C-D)** Stimulation of Gluc-LHCGRv4 with rec-hCG **(C)** or rec-LH **(D)** resulted in cAMP production similar to that observed for LHCGR^WT^-P2A-mCherry. On the contrary, in the case of other Gluc-LHCGR variants cAMP production was negligible. Data is representative of an experiment performed in triplicate and was repeated independently at least three times.

### Gluc functionality

To ensure that Gluc was in fact fused to the LHCGR on the membrane, as sometimes fused proteins might be cleaved by endogenous or exogenous proteases ^29^, we then analyzed the luminescence on cells and in the medium. As shown in **Figure 2B**, Gluc activity was virtually only in the cellular faction, whereas in the medium only an insignificant fraction of luminescence could be detected, likely from dead cells. Once we ensured that Gluc-LHCGR^WT^ was fully functional, we moved to analyze the ligands that will be used in BRET.

### Receptor activation

To ensure that the fusion to Gluc does not block the functionality of the chimeric LHCGR, we then determined their responsiveness to both hormones (rec-LH and rec-hCG) by measuring cAMP production. Stimulation of Gluc-LHCGRv4 with rec-hCG resulted in cAMP production comparable to that of the WT receptor, whereas stimulation of other Gluc-LHCGR architectures (v1, v2, v3) resulted in negligible cAMP production upon receptor stimulation (**Figure 2C**). Similar results were observed for cells expressing Gluc-LHCGR stimulated with rec-LH (**Figure 2D**).

### Biosynthesis of the hormone-eGFP fusions

We then look into the eGFP-fused hormones. The cellular localization of the eGFP-tagged hormones was assessed by confocal microscopy. In all cases, the acceptor proteins were not dispersed in the cytoplasm, but they were visible in secretion trafficking route (**Figure 3A**), as we have previously reported ^26^.

**Figure 3.**
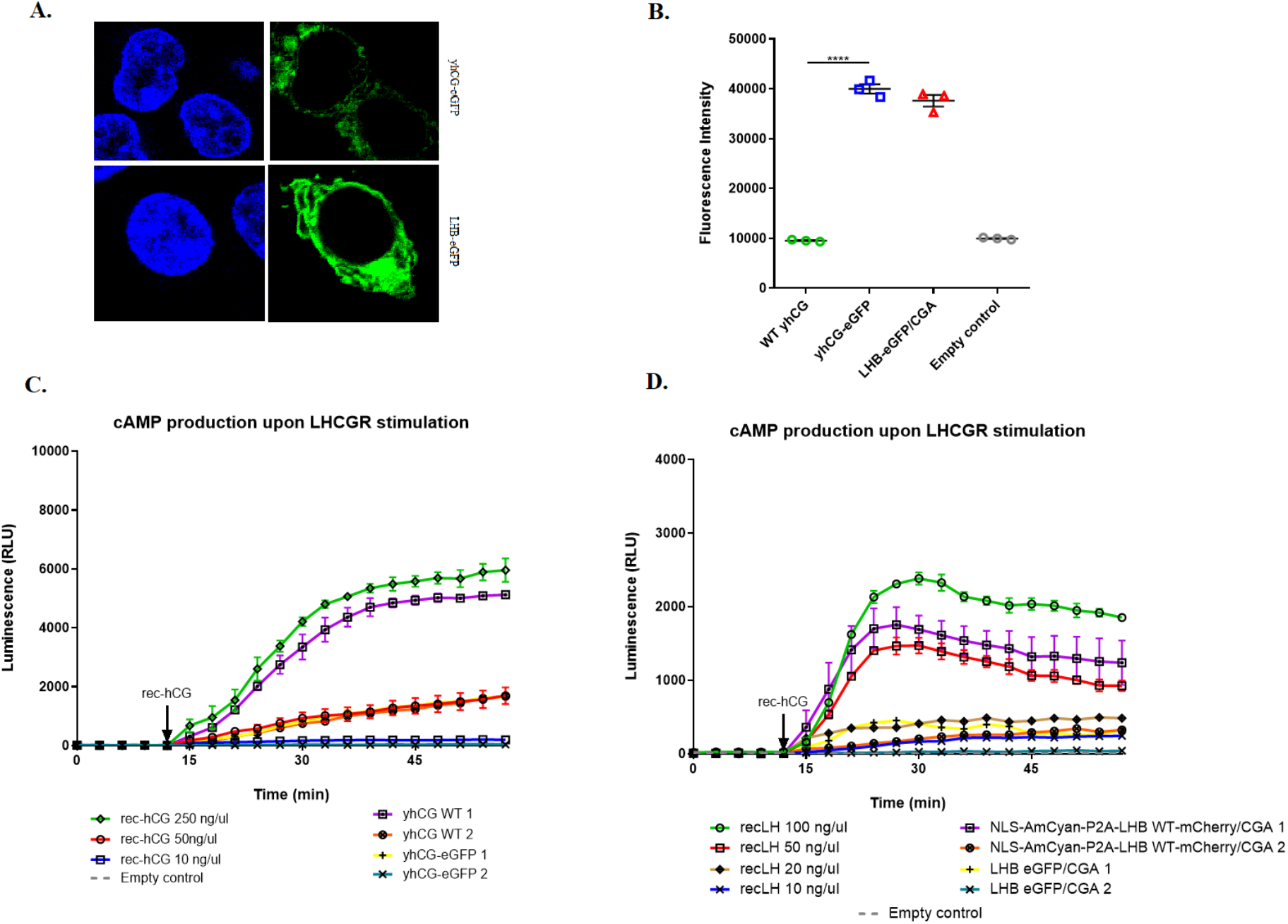
Cellular localization of acceptor moieties and cAMP generation upon receptor stimulation with eGFP-labeled ligands. **(A)** Confocal microscopy revealed the presence of both yhCG-eGFP and LHB-eGFP/CGA within the secretion route (ER/Golgi/vesicles). **(B)** Fluorescence intensity revealed that the fluorescence of medium collected from the cells transfected with yhCG-eGFP or LHB-eGFP/CGA were significantly higher as compared to the fluorescence of medium taken from the mock transfected control or untagged yhCG. The fluorescence intensity of LH-eGFP was lower than that of yhCG-eGFP which suggests a poorer secretion. Data is representative of an experiment performed in triplicate and was repeated independently at least three times. **** p<0.0001 **(C-D)** Both chimeric hormones, yhCG-eGFP and LH-eGFP (LHB-eGFP co-expressed with CGA) retained their ability to activate the LHCGR. **(C)** LHCGR stimulation with concentrated yhCG^WT^ resulted in receptor activation comparable to that induced by stimulation with rec-hCG at concentration of 250ng/μl. In turn, the use of concentrated eGFP-fused hCG resulted in LHCGR activation comparable to that observed for rec-hCG at concentration of 50ng/μl. **(D)** In turn, the use of a concentrated medium containing NLS-AmCyan-P2A-LHB^WT^-mCherry and CGA resulted in LHCGR activation similar to that induced by rec-LH at concentration of 50ng/μl, whereas receptor stimulation with concentrated eGFP-fused LH resulted in its activation at a level similar to that observed for the use of rec-LH at concentration of 20ng/μl. Data is representative of an experiment performed in triplicate and was repeated independently at least three times.

### Measurement of fluorescence and receptor activation with hormone-eGFP

To ensure that eGFP-fused hormones were secreted and able to activate their cognate receptor, we collected the media and analysed both florescence and by their ability to activate the WT LHCGR. We first analyzed if the chimeric hormones were properly secreted, to do this we measured eGFP fluorescence in medium collected from cells transfected with either yhCG^WT^, yhCG^WT^-eGFP, LHB^WT^-eGFP/CGA or mock plasmids, showing that the fluorescence in medium from yhCG^WT^-eGFP and LH^WT^-eGFP secreting cells was significantly higher than that of yhCG^WT^ to mock transfected cells (**Figure 3B)**. A clear difference in fluorescence between yhCG^WT^-eGFP and LH^WT^-eGFP could be noted, whereas the levels of LH^WT^-eGFP was less than yhCG^WT^-eGFP

We then tested whether the eGFP-tagged hormones retained their ability to stimulate the LHCGR. The concentration of hormone was equalized taking advantage of eGFP. In **Figure 3C** only the concentrations of the recombinant hormones which caused LHCGR activation to a level comparable to the activation caused by the use of either concentrated or non-concentrated WT or eGFP-coupled hormones are shown. The use of concentrated WT yhCG resulted in LHCGR activation comparable to that induced by the use of rec-hCG at concentration of 250ng/μl. In turn, the use of non-concentrated cell culture medium resulted in activation at a level comparable to that of rec-hCG at 50ng/μl, as well as for the concentrated yhCG-eGFP (**Figure 3C**). When using concentrated medium collected from cells co-transfected with NLS-AmCyan-P2A-LHB^WT^-mCherry and CGA, LHCGR activation was comparable to that observed for rec-LH at concentration of 50ng/μl. The stimulation with non-concentrated medium resulted in LHCGR activation comparable to that obtained for the rec-LH concentration of 10ng/μl. In the case of cells co-transfected with LHB-eGFP and CGA, the use of concentrated eGFP-fused LH resulted in receptor activation similar to that observed for rec-LH at concentration of 20ng/μl, while the use of non-concentrated eGFP-fused LH was too low to be measured (**Figure 3D**).

### BRET assay

Knowingly that the chimeric hormone (acceptor) and the donor Gluc-LHCGR function just as their WT counterparts, we further analyzed their interactions using BRET. The ratio between acceptor and donor emissions calculated for LHCGR stimulated with WT yhCG is comparable to the ratio obtained for the receptor stimulated with hormone-free medium.

By contrast, the acceptor:donor ratio calculated for cells expressing Gluc-LHCGR and stimulated with eGFP-fused yhCG is 2.9-fold higher as compared to the ratio of cells stimulated with WT yhCG (**Figure 4A).** In the case of LHCGR stimulation with eGFP-fused LH, the calculated ratio between the acceptor and donor was 3.4-fold higher than that calculated for the stimulation with NLS-AmCyan-P2A-LHB^WT^-mCherry/CGA which was negligible and comparable to the hormone-free medium (**Figure 4B**).

**Figure 4.**
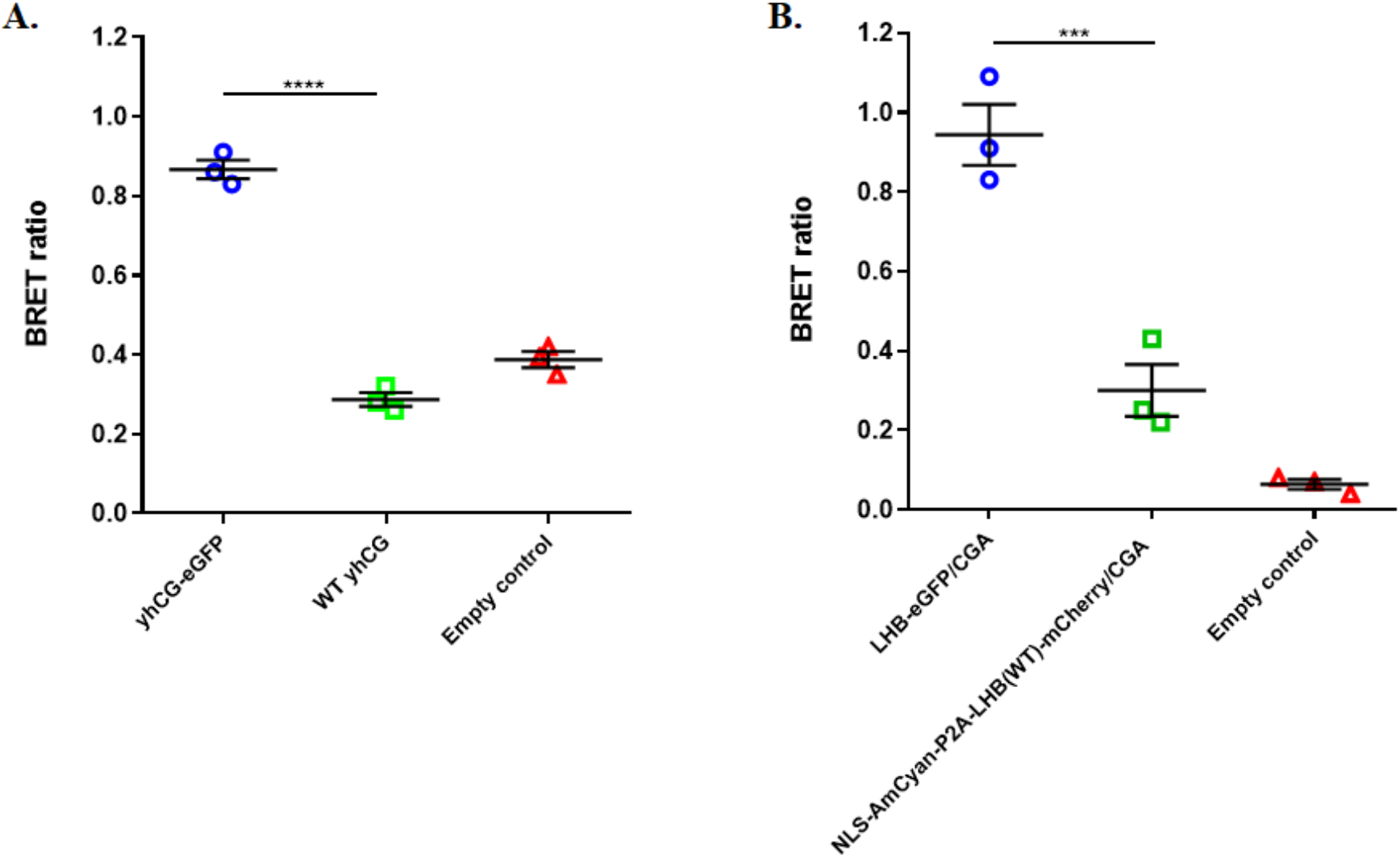
Measurement of BRET. **(A)** Stimulation of LHCGR with yhCG-eGFP resulted in an almost 3-fold higher acceptor:donor ratio as compared to the WT yhCG. BRET ratio calculated for the cells stimulated with WT yhCG was comparable to the BRET ratio calculated for hormone-free medium. **(B)** Calculation of acceptor:donor ratio showed that LHCGR stimulation with LH-eGFP resulted in an approximately 3.4-fold higher BRET ratio than in the case of stimulation with AmCyan-P2A-LHB^WT^-mCherry/CGA. Data is representative of an experiment performed in triplicate and was repeated independently at least three times. **** p<0.0001, *** p=0.0005

### Application of ligand-binding assay based on BRET

As described in the materials and methods section, a two-part experiment was performed to demonstrate the functionality of the ligand-binding assay. We selected the yhCG^Glu80Ala^ mutant, previously reported as yhCG^Glu56Ala^, to test our assay. At first, we noticed that the secretion of yhCG^Glu80Ala^-eGFP was lower than yhCG(WT)-eGFP, and thus, taking advantage of the eGFP, determine the cellular localization of mutant yhCG-eGFP using confocal microscope. Cellular imaging showed that secretion of mutant hormone was significantly reduced as compared to the yhCG-eGFP expression (**Figure 5A**).

**Figure 5.**
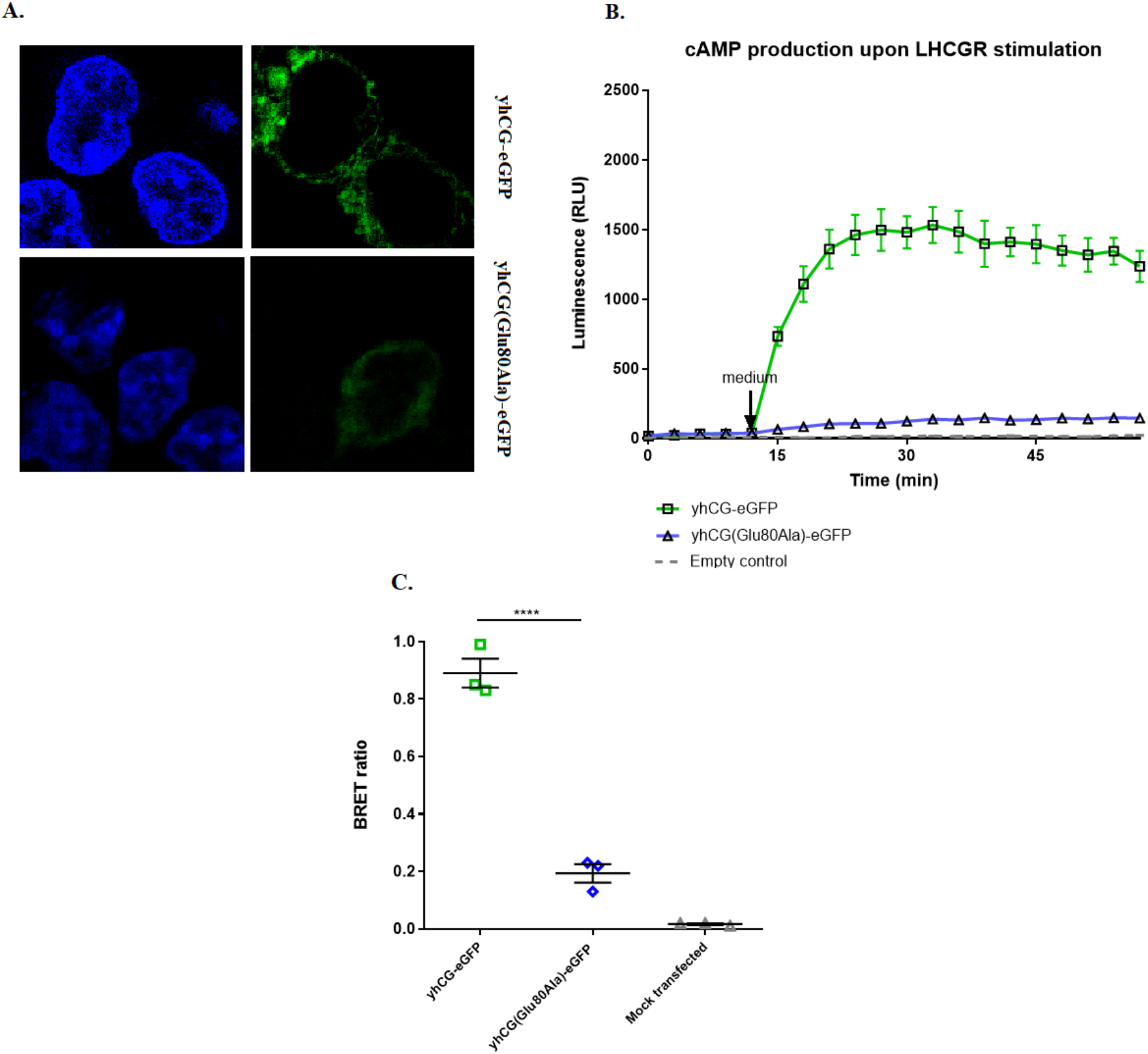
Molecular characterization of yhCG Glu80Ala mutation. **(A)** Confocal microscopy showed that the expression of yhCG^Glu80Ala^-eGFP was significantly decreased as compared to the yhCG^WT^-eGFP expression. **(B)** The results of GloSensor cAMP assay revealed that stimulation of GS-LHCGR with equimolar concentrations of yhCG^Glu80Ala^-eGFP resulted in 7.3-fold less cAMP production as compared to the yhCG^WT^-eGFP. **(C)** The acceptor:donor ratio obtained for the LHCGR stimulation with medium containing yhCG^Glu80Ala^-eGFP was 4.6-fold lower as compared to the BRET ratio calculated for the yhCG^WT^-eGFP. Data is representative of an experiment performed in triplicate and was repeated independently at least three times. **** p<0.0001

Then, we performed GloSensor Assay using equalized hormone concentrations which revealed 7.3-fold less cAMP production after yhCG^Glu80Ala^-eGFP stimulation as compared to yhCG^WT^-eGFP (**Figure 5B**).

Afterwards, BRET was performed using equalized hormone concentrations to demonstrate the interaction between the mutant ligand and LHCGR. The conducted experiment revealed that in the case of using a medium containing yhCG^Glu80Ala^-eGFP, the acceptor:donor ratio was 4.6-fold lower as compared to the BRET ratio calculated for the equimolar yhCG^WT^-eGFP moiety (**Figure 5C**).

Since hormone binding depends in both the hormone and the receptor, we then selected two mutations in the ECD of the LHCGR that have been previously described as a binding-deficient - Cys131Arg and Ile152Thr^21,22^ and one able to bind but unable to signal - Glu354Lys^23^. First, we tested whether these mutant receptors fused with Gluc, where then expressed on the plasma membrane by flow cytometry (**Figure 6A**)

**Figure 6.**
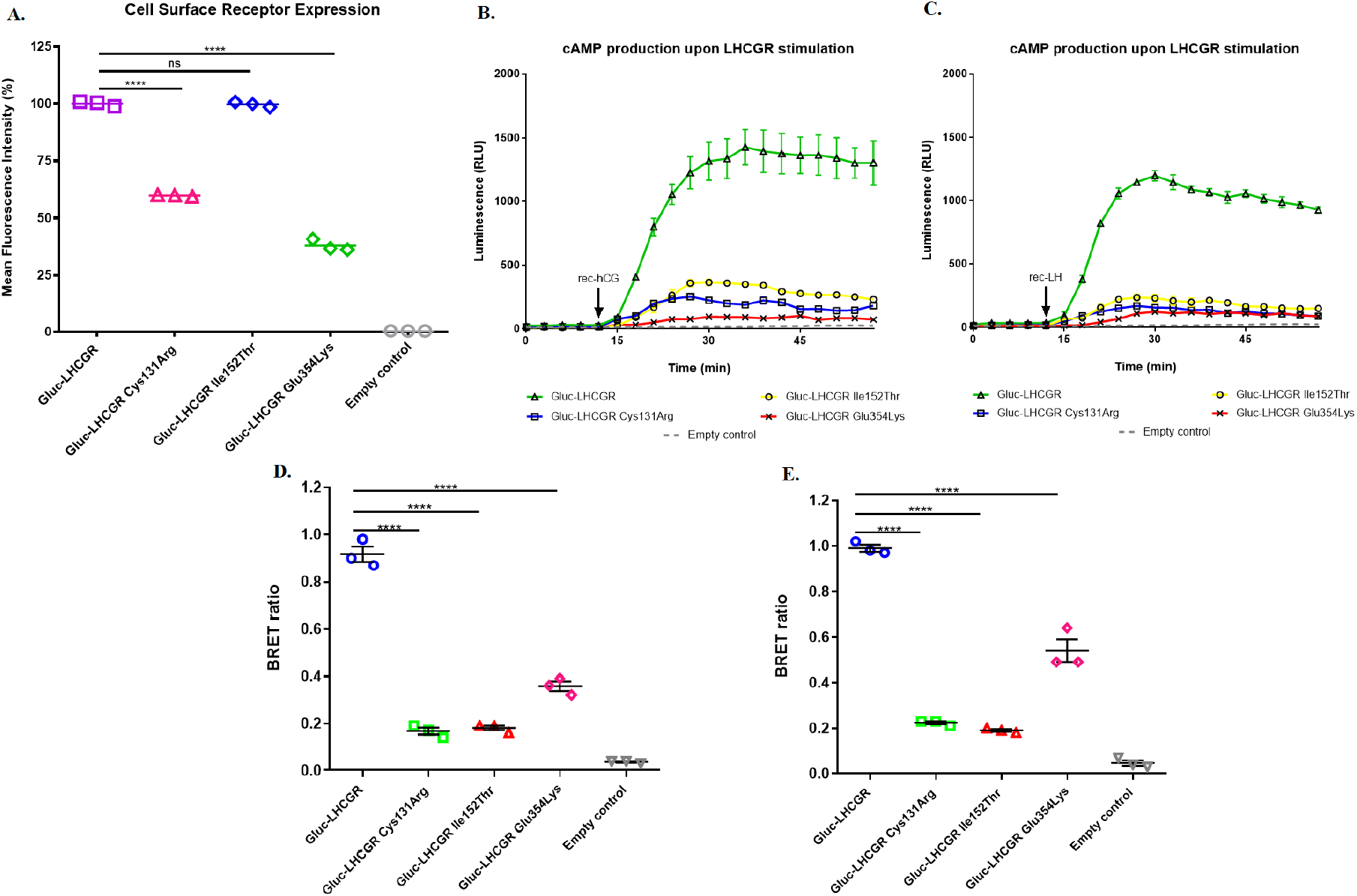
Molecular characterization of LHCGR mutants. **(A)** Flow cytometry analysis revealed that the membrane expression of Gluc-LHCGR^Cys131Arg^ was 1.7-fold lower, whereas the expression of Gluc-LHCGR^Ile152Ihr^ was almost the same as the membrane expression of Gluc-LHCGR^WT^. The lowest mean fluorescence intensity was observed for Gluc-LHCGR^Glu354Ala^ and it was 2.6-fold lower as compared to the Gluc-LHCGR^WT^. Data is expressed as MFI ± standard error of the mean (SEM) of three independent experiments. **** p<0.0001 **(B)** Activation of Gluc-LHCGR^Cys131Arg^ and Gluc-LHCGR^Ile152Thr^ with rec-hCG resulted in 7.4-fold and 4.9-fold lower cAMP production, respectively, as compared to Gluc-LHCGR^WT^. The lowest cAMP production was observed for the Gluc-LHCGR^Glu354Lys^ and it was 11-fold lower than that noted for Gluc-LHCGR^WT^. **(C)** Similar results were obtained with LHCGR stimulation with rec-LH. Here, 6.2- and 4.2-fold lower receptor activation was observed for Gluc-LHCGR^Cys131Arg^ and Gluc-LHCGR^Ile152Thr^, respectively. Similarly, the lowest cAMP production was observed for Gluc-LHCGR^Glu354Lys^, which was 14.4-fold lower as compared to Gluc-LHCGR^WT^. **(D)** BRET assay revealed that the acceptor:donor ratio obtained for the Gluc-LHCGR^Cys131Arg^ was 5.5-fold lower, whereas the BRET ratio calculated for Gluc-LHCGR^De152Thr^ was 5.1-fold lower as compared to the Gluc-LHCGR^WT^ stimulated with rec-hCG. On the contrary, acceptor:donor ratio calculated for the Gluc-LHCGR^Glu354Lys^ stimulated with rec-hCG was 2.6-fold lower as compared to the WT. **(E)** In the case of stimulation with rec-LH, similar values were obtained for the Gluc-LHCGR^Cys131Arg^ and Gluc-LHCGR^Ile152Thr^. The acceptor:donor ratio calculated for the Gluc-LHCGR^Cys131Arg^ was 4.4-fold lower and 5.2-fold lower for the latter mutant as compared to the Gluc-LHCGR^WT^. The BRET ratio obtained for the Gluc-LHCGR^Glu354Lys^ stimulated with the rec-LH was 1.8-fold lower as compared with the Gluc-LHCGR^WT^. Data is representative of an experiment performed in triplicate and was repeated independently at least three times. **** p<0.0001.

Once we knew that the mutant receptors are localized on the membrane, we tested if they are being activated by stimulation with either rec-LH or rec-yhCG and whether they bind to eGFP-fused ligands (**Figure 6B**).

As presented in **Figure 6**, all tested LHCGR mutants were activated as a result of stimulation with either rec-hCG **(Figure 6B)** or rec-LH **(Figure 6C).** Nevertheless, in the case of Gluc-LHCGR^Cys131Arg^ and Gluc-LHCGR^Ile152Thr^, several fold lower cAMP production was observed as compared to Gluc-LHCGR^WT^, whereas Gluc-LHCGR^Glu354Lys^ was distinguished by the lowest cAMP production. In turn, the BRET assay analysis **(Figures 6D and 6E)** revealed that the acceptor:donor ratios calculated for Gluc-LHCGR^Cys131Arg^ and Gluc-LHCGR^Ile152Thr^ were approximately 5-fold lower as compared to the ratio calculated for Gluc-LHCGR^WT^. On the contrary, the BRET ratio obtained for Gluc-LHCGR^Glu354Lys^ was 2.6-and 1.8-fold lower when stimulated with rec-hCG **(Figure 6D)** and rec-LH **(Figure 6E)**, respectively as compared to Gluc-LHCGR^WT^. These results corroborate pervious reports reporting that the decreased activation of LHCGR^Cys131Arg^ and LHCGR^Ile152Thr^ results from their decreased ability to bind the hormone and not from decreased expression on the plasma membrane. By contrast, the BRET values calculated for Gluc-LHCGR^Glu354Lys^ indicate that the decreased production of cAMP due to ligand stimulation is mainly due to decreased membrane expression of this receptor and its intracellular retention. In this case, the decrease in membrane expression of receptor correlates with a decreased BRET ratio. The obtained results are consistent with earlier reports on the above-mentioned mutations and confirm their influence on the LHCGR binding and signaling.

Finally, we tested whether our BRET (ligand:receptor binding) bioassay could be coupled to the cAMP (receptor activation) assay in the same samples, since the luminescence from Gluc and that of firefly luciferase (FFluc) are significantly different in light spectra as well as they require the use of different substrates ^14,29^. Since the readout of Gluc is immediate upon addition, while the cAMP GloSensor requires time stabilize and accumulate in cells, the assays were performed sequentially (**Figure 7A**). The charts in Figure 7 show that cAMP (GloSensor) (**Figure 7B**) and BRET (**Figure 7C**) could be analyzed in the very same cells in a sequential manner.

**Figure 7.**
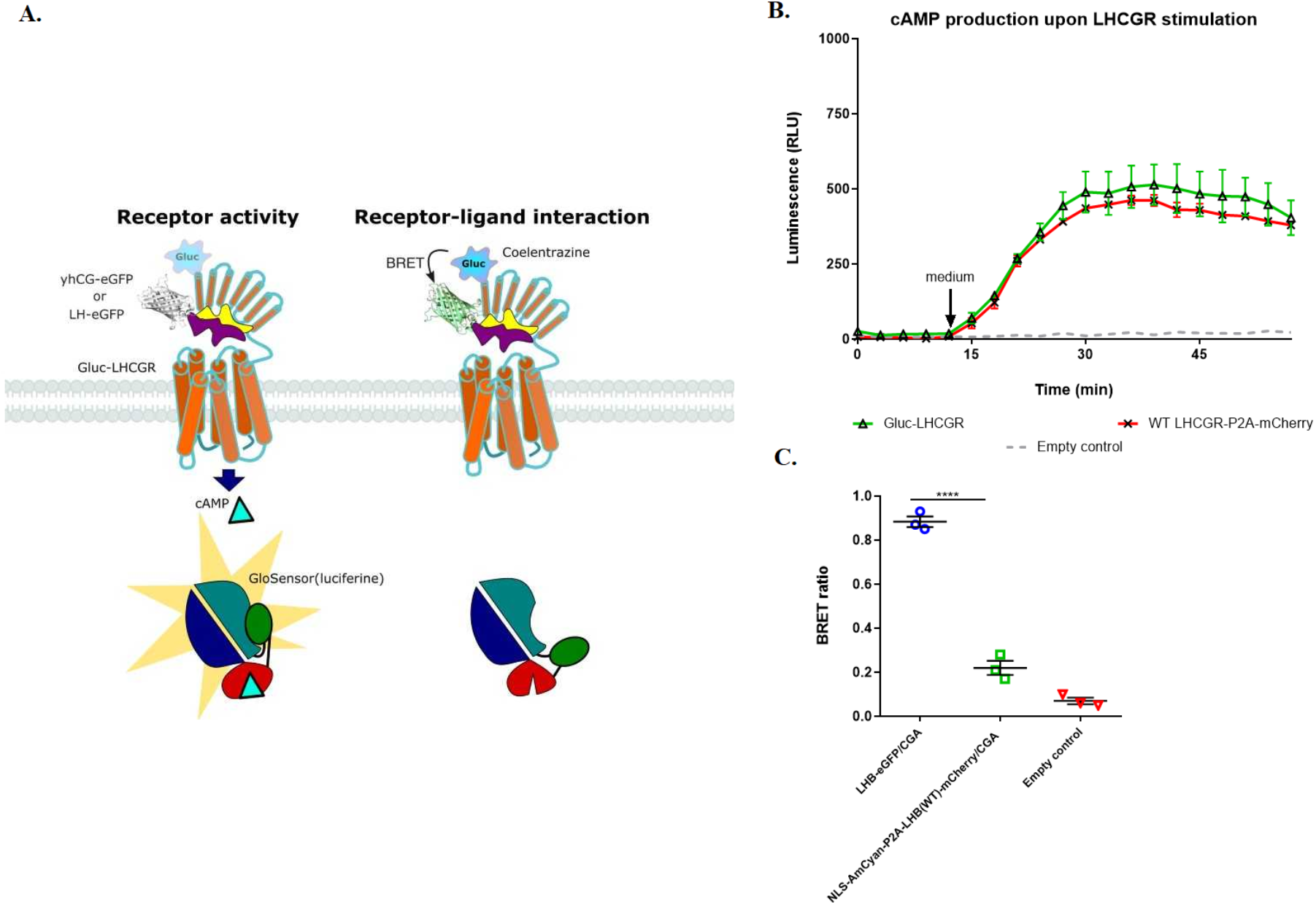
Coupling BRET assay with cAMP GloSensor assay. **(A)** Schematic representation of the parallel or sequential analyses. **(B)** Stimulation of Gluc-LHCGR with eGFP-fused LH resulted in cAMP production similar to that observed for LHCGR^WT^-P2A-mCherry. Data is representative of an experiment performed in triplicate and was repeated independently at least three times. **(C)** Calculation of BRET ratio revealed that Gluc-LHCGR stimulation with LH-eGFP resulted in 4-fold higher BRET ratio than in the case of stimulation with unlabeled hormone. Data is representative of an experiment performed in triplicate and was repeated independently at least three times. **** p<0.0001.

## Discussion

The methods used so far to study the interactions between ligands and their receptors were mainly based on the use of a radioisotope-labeled ligand associated with the need to ensure appropriate conditions in the laboratory and safety protocols. Usually such assays involve the use of isolated cell membranes ^31–33^ and competition is “cold” hormone to show specificity. Furthermore, it generates a significant cost of research resulting from the need to dispose of radioactivity waste. We depart from the traditional labeling of ligand with a radioisotope, replacing it with a fluorescent protein. Nevertheless, the mere replacement of the radioisotope with a fluorescent protein has already been used in FRET-based methods, where a second fusion protein consisting of an appropriate receptor and a second fluorescent protein is used ^11^. The use of FRET-based methods is distinguished by many advantages, such as high spatial and temporal resolution, but their application is associated with certain limitations due to the necessity to use an external light source to excite the donor protein. This in turn generates a high background, thus lower signal-to-noise ratio, and high heterogeneity between cells/samples. Furthermore, other significant disadvantages of FRET application in study of protein-protein interactions are donor photobleaching, which results in signal decrease over time, as well as the phenomenon of spectral overlap which leads to the bleed-through requiring subsequent corrections ^34^.

In this research, the BRET phenomenon was applied in ligand-binding assay in which luciferase catalyzes the oxidation of the substrate and its conversion to its derivative, resulting in the emission of a photon. For this purpose an acceptor fusion protein was designed and created, in which the extracellular domain of the LHCGR was fused not with a fluorescent protein, but with *Gaussia* luciferase (Gluc). Therefore, the BRET method is distinguished by the lack of background noise and thus higher sensitivity as compared to the FRET-based assay ^35^. Additionally, due to the fact that this method does not require excitement with an external light source, it constitutes a more accessible alternative to the FRET-based method due to simpler instrumentation requirements ^36^. Another advantage of BRET method application in the study of protein-protein interaction is the constant photon emission which enables the performance of studies over time. Furthermore, in contrast to the FRET, there is no false excitation or bleed-through phenomenon.

Hitherto, the BRET phenomenon has been used mainly to study the interactions between the GPCRs and to determine whether receptors form dimers or oligomers ^37,38^. The application of the BRET phenomenon in the study of receptor-ligand interactions may be of great biological importance in studies focused on binding of LHCGR with its corresponding ligands - hCG and LH. This method may constitute a useful tool for molecular characterization of novel and already identified mutations in the genes encoding both hormones’ subunits as well as their receptor.

The LHCGR is an extremely important constituent for the proper functioning of female and male reproductive systems through stimulation of ovulation in women and induction of testosterone production in men. Although mutations in the *LHCGR* gene are extremely rare they can have a huge impact on the sexual development and fertility of the patients affected by mutations ^39,40^. Their discovery and subsequent examination gives a better insight into the importance of LHCGR for human reproduction as well as they expand the current state of knowledge about the entire family of GPCRs. Mutations in genes encoding hormone-specific subunits are even more rare and only cases of inactivating mutations in these genes have been identified. Nonetheless, similarly to the mutations affecting *LHCGR* gene, these mutations are associated with a wide range of symptoms such as delayed puberty, hypogonadotropic hypogonadism and infertility in men as well as secondary amenorrhea and infertility in women^4^. Most often, aforementioned mutations result in impaired biosynthesis, abnormal posttranslational modification, incorrect heterodimerization with common subunit, impaired secretion of the hormone and its binding with receptor ^41^. Moreover, we show that binding- and activation-assays can be run on the same samples, what saves time and money, and generates data with enhanced kinetics. In summary, the BRET-based ligand-binding assay developed by us enables to investigate the effect of mutations identified in both the *LHCGR* gene as well as in genes encoding its ligands’ subunits. Additionally, it is possible to apply this method in the study of mutations present in genes encoding other GPCRs, and in particular glycoprotein receptors, and their ligands after appropriately introduced changes using genetic engineering. A significant advantage distinguishing this method is the use of live cells instead of isolated cell membranes. This in turn is associated with another advantage of the BRET-based method, which is the possibility of simultaneous use of two different luciferases - *Gaussia* and *Firefly*. This is possible due to the use of different substrates for these enzymes and due to their different emission spectra and thus the use of different filters set for the signal measurement. Therefore, it is possible to test ligand-binding using *Gaussia* luciferase as well as to investigate receptor activation and downstream signaling pathways using *Firefly* luciferase in the same cells. Simultaneous measurement of ligand-binding assay and receptor activation is possible, nonetheless limited by the possession of the appropriate equipment enabling the separation of the emission spectra for both luciferases and the fluorescent protein. Furthermore, another advantage of this BRET-based assay is the possibility to measure the membrane expression of LHCGR, or any other GPCR, by adding coelenterazine to the cells expressing the Gluc-fused receptor, followed by the luminescence readout. The level of observed luminescence correlates with the level of relative membrane expression of the GPCR under study as described by Rodríguez et al. on the example of Gluc fused to the extracellular part of the Cannabinoid receptor 1 (CB1) ^29^.

To conclude, the method of studying the interaction between the glycoprotein and the glycoprotein receptor described by us is a simpler and faster alternative to the ligand-binding methods used so far, at the same time enabling the assessment of receptor membrane expression as well as the study of receptor activation using downstream signaling assays.

## Acknowledgements

Authors would like to thank Addgene, and the colleagues for the support to the greater scientific community.

